# Lack of Synergistic Outcomes with Roflumilast Combined with Levetiracetam or Adipose Stem Cell Secretome After Spinal Cord Injury

**DOI:** 10.1101/2024.07.02.601664

**Authors:** Carla Sousa, Rui Lima, Eduardo D. Gomes, Deolinda Silva, Jorge Cibrão, Tiffany Pinho, Diogo Jorge, João Afonso, Joana Martins-Macedo, Andreia Monteiro, António J. Salgado, Nuno A. Silva

## Abstract

The Spinal Cord Injury (SCI) pathophysiology is highly complex, contributing to a poor prognosis and lack of effective treatments. Previously, we demonstrated that Roflumilast (Rof), leads to functional recovery when tested in a SCI contusion model. However, it is unlikely that Rof treatment on its own could fully restore the spinal cord. Therefore, our objective was to scrutinize the synergistic effects of combining Rof with neuroprotective approaches. Herein we tested two therapies, firstly, Rof combined with Levetiracetam (Lev), and in a second phase, the complementary interplay between Rof and Adipose Stem Cells secretome (Sec). We induced SCI using a weight drop trauma model at the T8 level. Functional recovery was assessed according to the Basso, Beattie, and Bresnahan scale, Activity Box Test, Motor Swimming Test, and Von Frei test. Results indicate that the unilateral use of Rof, Lev, or Sec was effective in promoting functional recovery. However, the combination of Rof + Lev or Rof + Sec did not lead to an improvement in functional outcomes when compared to standalone treatments. Moreover, the combination of Rof + Sec actually led to worst functional outcome than the single treatments. Further studies are needed to find a combinatorial treatment that can lead to superior therapeutic effects with potential clinical application.

## Introduction

Traumatic spinal cord injury (SCI) can result in dramatic neurological disability with a strong social and psychological impact. Primary injury is caused by mechanical forces applied directly to the spinal cord at the time of injury [1]. After that, a cascade of secondary events take place with a progressive process of post-traumatic damage [2], which includes permeability and vascular variations, free radical formation and lipid peroxidation, ionic disruption and glutamate excitotoxicity, dysfunctional inflammatory reaction, apoptosis, and finally glial scarring. The events of the secondary injury create an inhibitory microenvironment that blocks axonal regeneration, greatly contributing to the failure of tissue repair observed after SCI. The identification of inhibitory proteins in central nervous system (CNS) myelin (such as MAG, Nogo, and OMgp) [3–7], greatly contributed to our understanding of the molecular basis of axonal regeneration. For instance, it was discovered a correlation between neuronal cAMP levels and inhibition of neurite outgrowth by myelin debris. When levels of endogenous cAMP were kept high, a significant neurite outgrowth could be observed, even under the presence of myelin [8]. For that reason, increasing the neuronal cAMP levels is known as one of the possible targets to promote axonal regeneration. For example, it has been shown that inhibiting the action of Phosphodiesterase-4 (PDE4), an enzyme responsible for the hydrolysis of cAMP, increases the levels of cAMP, which activates protein kinase A (PKA) and successively CREB, thereby promoting transcription of genes associated with synaptic plasticity, axonal growth, and neurogenesis [9, 10]. Recently, our team tested an FDA-approved PDE4 inhibitor, Roflumilast (Rof), as a strategy to promote axonal regeneration and repair after SCI. We observed that Rof was able to increase axonal regeneration in vitro and lead to functional recovery in vivo. These results were accompanied by an increase in the cAMP levels in the spinal cord tissue as well as a significant decrease in the cavity size and less reactive microglia cells [11].

Together with axonal growth inhibition, one of the central pieces of the SCI pathological process is glutamate excitotoxicity. After the SCI, calcium (Ca2+) flux at the synaptic cleft regulates the release of glutamate. As glutamate receptors (NMDA and AMPA) become overactivated, sodium (Na+) and Ca2+ influx increases. As a result of this ionic dysregulation, neuronal and glial cells, especially oligodendrocytes and neurons, become vulnerable to cell death [12, 13]. Moreover, axonal degeneration is mediated by Ca2+ influx from the endoplasmic reticulum (ER) via the IP3 receptor that promotes mitochondrial permeability [14]. Globally, glutamate concentrations increase to neurotoxic levels (excitotoxicity) resulting in axonal demyelination and neuronal cell death [15–17]. Thus, modulating excitatory amino acid levels may be an effective strategy to promote neuroprotection after SCI trauma. Several drugs that modulate excitotoxicity have been tested over the years. Our team has recently shown that Levetiracetam (Lev) had a neuroprotective role after SCI. Lev is an antiepileptic drug that suppresses presynaptic neurotransmitter release by binding to synaptic vesicle protein SV2A [18–21]. Using a cervical and thoracic rat SCI model, our team demonstrated that Lev treatment promotes significant improvements in both gross and fine motor function, revealing a significant decrease in cavity contusion size, as well as higher neuronal and oligodendrocyte survival [22].

Another promising therapeutic approach that our team has been testing is the use of the proteins and extracellular vesicles secreted by Mesenchymal Stem Cells, namely the ones derived from adipose tissue (ASCs). Different studies point to the secreted factors and vesicles, known as the secretome, released by these cells as agents for the regenerative effect rather than ASCs differentiation. The secretome can be defined as the factors that are secreted by a cell, tissue, or organism to the extracellular space under defined time and conditions [23]. We have previously reported that the ASC secretome is an inductor of vascularization [24], as well as a promoter of neurite growth, leading to functional recovery when injected in mice with thoracic contusive or full transection SCI [25].

While individual therapies have shown promise in preclinical models, the fact is that they have been unable to demonstrate clinical efficacy when tested in humans. For this reason, it is important to search for complementary treatment strategies that can address diverse aspects of SCI pathology and produce higher levels of regeneration. In this sense, herein we combined the neuroprotective effects of Lev with the neuroregenerative effects of Rof. Moreover, we also searched for synergistic or additive therapeutic effects between the Rof and ASCs secretome.

## 2. Results

### 2.1. Roflumilast and Levetiracetam alone or combined treatments promote functional recovery after SCI

Four hours post-injury, the animals were administered Lev (750 mg/kg) via IP injection. Subsequently, they received 3 intrathecal injections of Rof (100 ug/kg) at 2, 4, and 6 dpi. Following this, they were administered a weekly IP injection of Roflumilast (1 mg/kg) commencing from the second-week post-injury. The BBB test was performed to evaluate locomotor recovery in an open arena with transparent acrylic walls, 3 dpi and thereafter weekly for 7 weeks. Results revealed that although all groups presented a spontaneous recovery over time, only treated animals showed a significant motor recovery when compared to saline (Figure 1A, p = 0.0064). However, Tukey’s multiple comparisons test did not reveal any difference amongst treatments when administered alone or in combination. At 8 weeks post-injury, animals treated with Rof (6.914 ± 3.281), Lev (7.654 ± 3.350), or Lev + Rof (7.264 ± 3.627) were able to extensively move the three joints of the hindlimb, while animals treated with saline (5.222 ± 2.665), only achieved slight movement of two joints and extensive movement of the third. The locomotor analysis considering two significant indicators of functional recovery, namely, body weight support and coordination, revealed significant differences when analyzed over time between drug-treated animals and the saline-treated group. Only 20% of the animals treated with saline recovered body weight support contrasting with 44% in the Lev (p=0.0001), 45% in the Rof (p<0.0001), and 62,5% in the Lev+Rof treated animals (p<0.0001; Figure 1B). Concerning coordination, none of the saline-treated animals recovered any level of coordination, while 33% of Lev, 45% of Rof, and 50% of Lev+Rof treated animals recovered at least occasional coordination (Figure 1C). Despite no statistical synergistic or additive effects were detected between the drugs, the animals treated with both drugs presented a higher percentage of recovery in body weight support and coordination.

**Figure 1.**
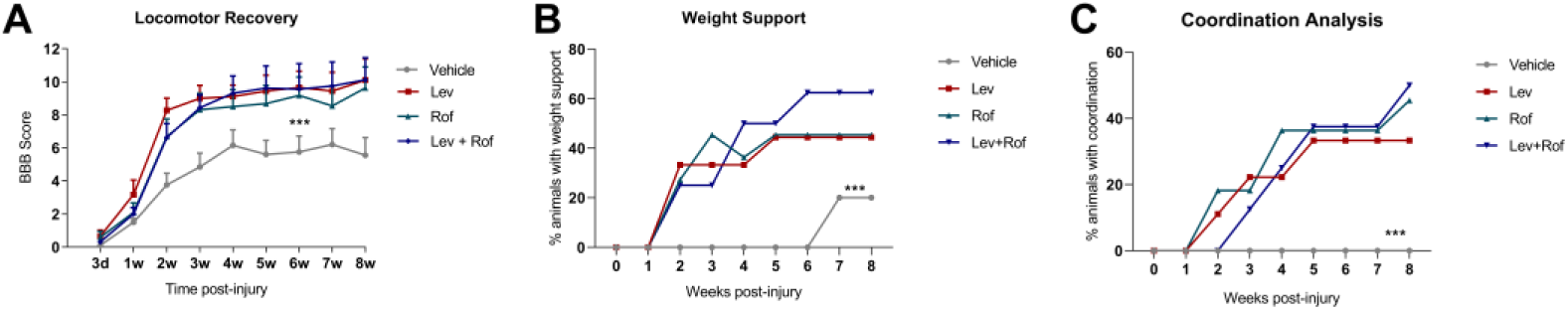
Evaluation of functional recovery after spinal cord injury in animals treated with Rof, Lev, Rof+Lev or vehicle. A) Locomotor recovery analysis demonstrated that only treated animals presented significant motor recovery when compared to vehicle. No differences were observed between the treatments when administered alone or in combination. B) Weight support analysis demonstrated that animals treated with Lev, Rof, and Lev+Rof presented higher recovery body weight support contrasting with the vehicle-treated group. C) Coordination analysis demonstrated that vehicle-treated animals did not recover any level of coordination. All treatment groups presented significant improvement in coordination when compared to controls, however, no statistical differences were observed between treatments. Data was analyzed using 2-way ANOVA p=0.0064 followed by Tukey’s multiple comparisons test. Vehicle n = 10; Lev n = 9; Rof n = 11; Lev+Rof n = 8. Values shown as mean ± SEM. ***p<0.001.

The activity box test was used to assess the exploratory behavior of the animals by measuring the total distance traveled, and velocity. The test was done in an open arena with transparent acrylic walls. Distance and locomotor velocity were evaluated at 8 weeks post-injury. The animals of all groups roamed a similar distance with no statistical difference (Figure 2A, p= 0.0673). However, animals that received Lev, Rof, or a combination of both roamed with significantly more velocity than saline-treated animals (Figure 2B, p=0.0008). Interestingly, the combination of Lev + Rof had a statistically significant difference when compared to the Lev-treated group (p=0.0206).

**Figure 2.**
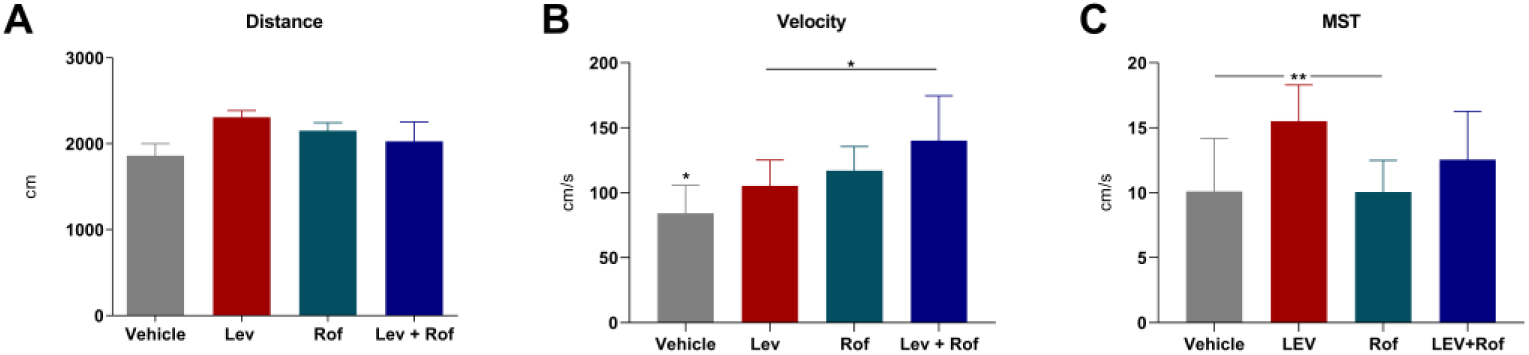
Analysis of exploratory behavior using the Activity Box Test. A) No differences were observed between all groups when analyzing total overground distance. B) Animals treated with our therapeutic agents presented higher velocity than vehicle-treated animals. A synergistic effect between Lev+Rof was observed but only when compared to the Lev-treated group. C) Animals treated with Lev showed a statistically significant difference when compared to the Rof and vehicle-treated groups on the motor swimming test (MST). Values shown as mean ± SEM. *p<0.05; **p<0.01.

The locomotor velocity during swimming was also analyzed. The test was performed on a quadrangular pool, where the rat had to reach a platform to get out of the water. All the trials were recorded by a video camera, after two days of training. In this analysis the Lev-treated animals showed a statistically significant difference when compared to the Saline and Rof groups. These animals presented higher velocity in water (Figure 2C, p=0.0015), however, there were no differences observed in Lev+Rof treated animals.

### 2.2. Roflumilast combined with ASCs secretome does not promote functional recovery after SCI

In the first 48h post-injury, the animals were administered 3 times with ASCs secretome (500ul/each time) via IV injection. Subsequently, they were administered a weekly IP injection of Rof (1 mg/kg) commencing from the second-week post-injury. The BBB was performed to evaluate locomotor recovery in an open arena, 3 dpi and thereafter weekly for 7 weeks (Figure 3A). Animals treated with the combination of Rof and Sec had a significantly worse performance than animals treated with Rof (p=0.0335) or Sec alone (p=0.0028). There were no statistical differences between the combinatorial group and the vehicle-treated animals (p= 0.3900). Moreover, there were no differences between Rof and Sec-treated animals (p= 0.9628).

**Figure 3.**
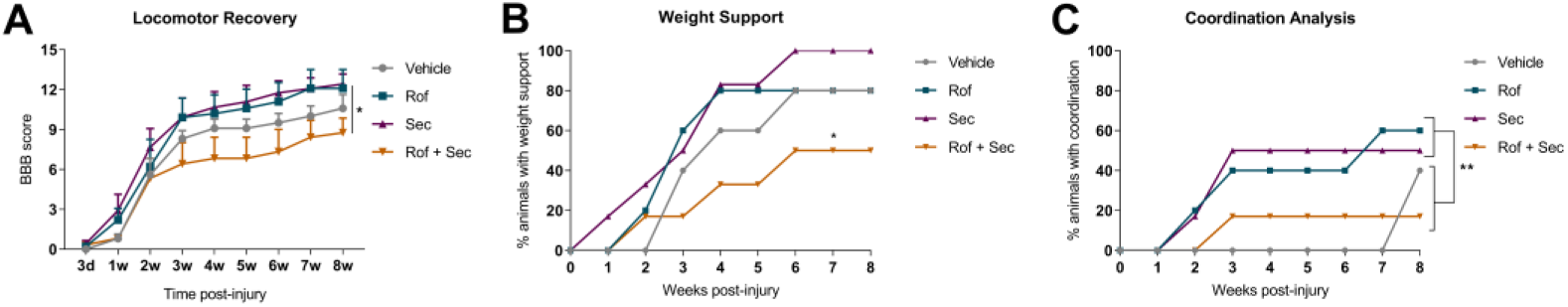
Evaluation of functional recovery after treatment with Rof, Sec, Rof+Sec, or vehicle. A) Locomotor recovery analysis demonstrated that the combination of Rof and Sec leads to a significant worse locomotor recovery when compared to animals treated with Rof or Secretome alone. B) Body weight support analysis revealed statistical differences between treatments with the combinatory therapy showing again detrimental effects. C) Coordination analysis revealed that when treatments are administered alone significant recovery can be achieved when compared with Rof+Sec or vehicle treatment. Differences over time were analyzed using 2-way ANOVA followed by Tukey’s multiple comparisons test. Vehicle n=5; Rof n=5; Sec n=6 and Rof+Sec n = 6. Values shown as mean ± SEM. *p<0.05; **p<0.01.

The locomotor analysis considering two significant indicators of functional recovery - body weight support and coordination - revealed significant differences between groups (Figure 3B and 3C). 50% of the animals treated with the combinatory approach recovery body weight support contrasting with 100% in the Sec (p <0.0001), and 80% in both the Rof (p=0.0002) and vehicle-treated animals (p= 0.0155). Concerning coordination, only 17% of combinatory-treated animals recovered at least occasional coordination, contrasting with 40% in the vehicle-treated group, however, no significant differences were detected between these groups (p= 0.6080). On the other hand, the animals treated with Sec or Rof alone significantly recovered more levels of coordination when compared with either Rof+Sec or vehicle-treated animals. Namely, 50% of Sec (p= 0.0013), and 60% of Rof-treated animals (p= 0.003) recovered at least occasional coordination (p values shown for the analysis against vehicle group). These results indicate the detrimental effects of the combinatorial treatment, however, when administered alone both compounds showed a therapeutic effect.

### 2.3. Treatments administered alone or in combination did not affect mechanical allodynia

The Von Frey test was performed at the seventh week post-injury to assess mechanical allodynia and hyperalgesia in the animals. This test aims to measure tactile sensitivity and pain responses and involves applying varying degrees of pressure to the paws of SCI animals, using filaments with different force intensities. Results did not reveal any statistical difference between all the treatments administered. Animals from all groups presented a low threshold to mechanical forces, and the treatment with Rof, Sec, or the combination of both did not improve or deteriorate mechanical allodynia in both the left and the right paws (Figure 4).

**Figure 4.**
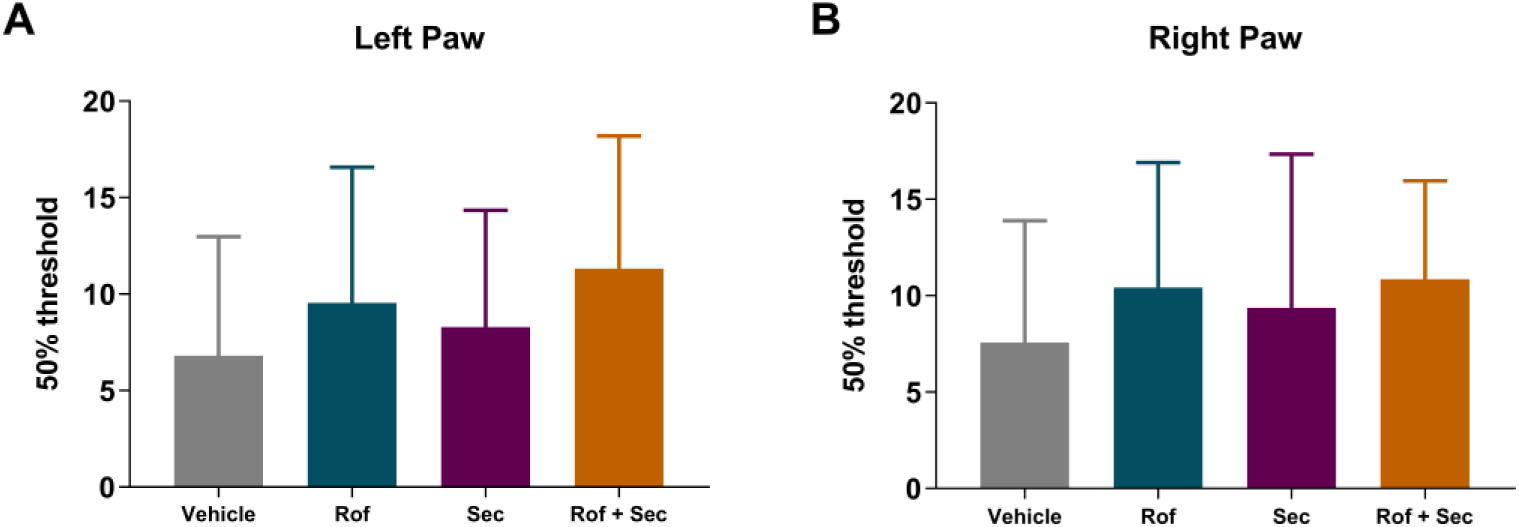
Results from Frey filaments applied both on the A) Left paw and B) Right paw. Statistical analysis did not reveal any differences between groups. Normality analysis using the Kolmogorov-Smirnov test revealed that data does not follow a Gaussian distribution, for this reason, the non-parametric Kruskal-Wallis test was performed followed by the Dunn’s multiple comparisons test. Vehicle = 5; Rof = 5; Sec = 6; Rof + Sec = 6. Values shown as mean ± SEM.

## 3. Discussion

Spinal cord regeneration is a complex physiological process, and no optimal treatment has yet been developed to fully repair the spinal cord after injury. There have been numerous studies in recent years showing the effects of several treatment options on SCI. Approaches such as biomolecules, stem cells, and 3D scaffolds are being explored to regenerate the spinal cord [24, 26–29]. Despite evidence suggesting that these strategies can be useful for spinal cord reconstruction, the fact is that single-based therapies have not been able to adequately improve clinical outcomes. Significant functional gains for individuals with SCI may require a combinatorial approach that addresses the multifactorial nature of SCI pathophysiology. In this sense, advances in combinatorial therapies, which can address different molecular and cellular secondary events of SCI, offer hope for spinal cord regeneration. Having this in mind, in this study we tested two different combinatorial approaches. Both aimed to promote neuroprotection in the acute phase and neuroregeneration in the chronic phase.

Initially, we aimed to evaluate the potential of a combined treatment approach using Lev and Rof. In our findings, we observed that individually, both Lev and Rof treatment can improve the functional outcome of SCI rats. These data reinforce our previous results where we demonstrated that Lev could protect the injured spinal cord in the acute phase leading to functional and histological recovery [22], and Rof can promote axonal regeneration in the chronic phase, leading as well to functional improvements [11]. The combined administration of Lev and Rof also significantly improved the motor function of the animals when compared with vehicle-treated animals. However, despite these statistical differences, we did not observe strong synergistic or additive effects resulting from our combination approach. In fact, in the BBB motor score, the animals treated with the agents alone or in combination do not differ. Nonetheless, it’s essential to highlight that we observed a higher overground velocity in the group that received the combined treatment when compared with Lev alone, while all treatments were also significantly better than the vehicle group. These may indicate an additive effect and, consequently, this combination may display promise for yielding better functional outcomes when administered together for SCI injuries. Further investigations focusing on various administration regimens, different dosage ratios, and alternative routes of administration are necessary to comprehensively understand whether these two drugs, when combined, might offer a higher degree of recovery compared to when administered individually. It is also important to point out that there was no observable distinction in velocity between the saline group and the Lev+Rof combination group during the swimming task. Moreover, only the Lev-treated animals presented significant improvements in this test. There are several similarities between swimming and walking, including the phasic relationship between the right and left limb flexion and extension, however, during swimming there are very low levels of both proprioceptive (loading of the limbs) and cutaneous feedback compared to the levels experienced during walking [30]. Cutaneous feedback plays a critical role in hindlimb locomotion, and it was demonstrated that animals exposed to phasic cutaneous feedback during swimming showed improved hindlimb function compared to those without this input [30]. The lack of environmental exploration during swimming, a behavior commonly observed during walking, might also have an impact on the assessment of some animals [31]. These differences between overground walking and swimming could explain our results. Notably, our previous publications indicated that Lev enhanced swimming recovery [22], while the same effect was not seen with Rof treatment [11]. Thus, it is important to direct additional experiments toward comprehending whether Lev distinctly enhances proprioception, particularly through the protection of the spinocerebellar tract and the dorsal columns, which are the two major pathways conveying proprioceptive information.

Given the lack of obvious additive effects between Rof and Lev, in the second part of the study, we aimed to assess whether Rof could potentially exhibit synergistic effects when combined with the secretome of ASCs. In the past years, our team has been investigating the therapeutic potential of the secretome derived from stem cells, particularly those displaying a mesenchymal phenotype, for regenerative applications in the CNS. The use of secretome offers numerous advantages over conventional stem cell transplantation, particularly in terms of manufacturing, storage, and practical application as a readily deployable biologic product. Notably, the secretome can be stored for extended periods without compromising its potency and quality [32, 33]. Large-scale production can be achieved under controlled laboratory conditions, and the biological product can be customized to desired cell-specific effects [34]. Importantly, secretome derivatives could circumvent potential challenges linked with cell transplantation, including limitations in cell availability, post-transplantation cell survival, immune compatibility, tumorigenicity, and risk of infection transmission.

Previous research has shown that the secretome of MSCs comprises essential proteins, growth factors, chemokines, and cytokines that facilitate the regenerative processes [35]. Various factors such as Nerve growth factor (NGF), Vascular endothelial growth factor (VEGF), Hepatocyte growth factor (HGF), Insulin-like growth factor 1 (IGF-1), Transforming growth factor-beta 1 (TGF-β1), Interleukin (IL)-10, Glial-derived neurotrophic growth factor (GDNF), basic Fibroblast growth factor (bFGF), Pigment epithelium-derived factor (PEDF), Cadherin 2 (CADH2), Semaphorin 7A (SEM7A), and Glial-derived nexin (GDN) have been identified as contributors to the neurotrophic and immunomodulatory effects of ASCs [36–38]. These factors have been associated with various critical processes such as neurogenesis, neuronal differentiation, proliferation [36–38], axonal growth, and migration [39–41]. Among the different sources available for isolating MSCs, adipose tissue-derived (ASCs) have emerged as particularly promising for SCI applications. Our previous investigations revealed that ASC secretome significantly promoted the neuronal differentiation of human neural progenitor cells and the axonal growth of dorsal root ganglion explants in vitro [42]. Notably, when compared with secretome derived from other sources like bone marrow or umbilical cord tissue, the ASC-derived secretome demonstrated superiority in these aspects. These outcomes are closely linked to the composition of ASC secretome, containing various well-recognized CNS-related neuroregulatory factors, including PEDF, Semaphorins (SEM), Cadherins (CDH), Interleukin (IL)-6, GDN, Clusterin (CLUS), Decorin (DCN), and Beta-1,4-galactosyltransferase 1 [43]. Moreover, our pre-clinical findings upon injecting the secretome following SCI have consistently shown a notable modulation of the immune response within the spinal cord tissue [25, 44].

Considering the well-documented beneficial effects of ASCs and our prior positive outcomes with Rof, herein we aimed to investigate the therapeutic potential of a combination involving the Sec and Rof. Surprisingly, our findings did not unveil any synergistic or additive effects between the two therapeutic agents. Unexpectedly, the combination of Sec and Rof led to detrimental effects on the motor function of the treated animals, both when analyzed on the BBB scale and when assessing the ability of the animals to regain body weight support or coordination. Despite these negative effects from the combined approach, our results reinforce the individual beneficial effects of the therapeutic agents when administered alone, which is consistent with our previous published data [25, 44].

Understanding the intricacies of combined treatments is complex. Investigation into the exact reasons for the detrimental effects in this specific combination would need to involve further research, considering various dosages, administration sequences, and the specific physiological response within the context of SCI models. To avoid unintended interaction between the several molecules of the Sec with the Rof, we administered the two therapeutic agents in two different time windows. The Sec was injected up to 48h post-injury and then the first injection of Rof happened only two weeks post-injury. In this sense, it is unlikely that a direct interaction between the proteins of the secretome and the Rof could have led to the observed detrimental results. However, we cannot discard some indirect interactions. It is possible that the two therapeutic agents modulate incompatible pathways, ASC secretome and Rof might target antagonistic pathways interfering with the actions of each other, leading to a compromised overall therapeutic response. Moreover, and despite the different time windows of administration used, the sequencing of administration might also play a role. A particular order between the two agents might be essential for their effectiveness, and the combination could have deviated from this optimal pattern. Based on all our results we discourage the combination of ASCs secretome and Rof for SCI repair.

Even though it is assumed by the scientific community that an effective treatment for SCI may need to rely on combinatorial treatments that target several hallmarks of SCI in an integrated manner, rather than developing an isolated approach, the present work demonstrated that testing combinatorial therapeutic approaches presents numerous challenges. An important challenge is the study of the interactions between therapeutic agents. Combinatorial approaches involve combining agents with different mechanisms of action with potential interactions. Understanding how these treatments interact is complex and may require extensive investigation. Determining the right combinations, doses, and administration schedules is also challenging. Each agent might have distinct pharmacokinetics and dynamics, and determining the most effective and safe dosing regimen will require careful study. Moreover, understanding the individual mechanisms of action and the potential overlap or divergence among multiple therapeutic agents is essential. However, even with this knowledge, it is difficult to predict whether the combined effects of therapeutic agents will be synergistic, additive, or antagonistic. These effects can be unpredictable and may not align with expectations based on the individual actions of the agents. Moreover, the translation to clinical settings will also be a big challenge for combined therapies. Translating single therapies for humans is proving to be complicated, thus doing it for combinatorial therapies will be even more complex. The design of clinical trial will have to consider the statistical power needed to detect treatment effects between single and combined agents, this may reveal to be cost-prohibitive and pose significant logistical challenges. Finally, obtaining regulatory approval for combinatorial therapies can be challenging due to the need for adequate safety and efficacy data for each agent and their combinations, however, it may be facilitated if the therapeutic agents to be combined are already FDA approved, as is the case of Lev and Rof.

The complexity of the SCI and the multifaceted nature of potential treatments will always be an issue, however, this field can also learn valuable lessons from the cancer field. Researchers working on cancer have made significant strides in understanding the challenges and benefits of combining different drugs for more effective treatments as well as on how to overcome some of the obstacles described above. Indeed, many researchers in the SCI field have been focused on finding the best combined treatment. In 2013, Martin Swab’s group tested the combination of two promising agents, the Anti-Nogo-A antibody and the ChABC enzyme. The combination treatment was more effective, leading to greater levels of neuronal sprouting and axon regeneration [45]. Another combination tested was the methylprednisolone and the Granulocyte colony-stimulating factor G-CSF. This combination promoted functional and histological improvements superior to those achieved by either of these drugs alone, demonstrating a synergistic effect [46]. Recently, Michael Fehling’s lab published a systematic meta-analysis that demonstrated that a combined application of scaffolds and MSCs is more effective than scaffolds and MSCs alone in improving motor function following SCI in animal models when used in the acute phase of injury [47]. Finally, Rilozule and Magnesium were also combined to protect the spinal cord against glutamate excitotoxicity. However, when tested together it was not observed any synergistic or cumulative effect, Riluzole was the only treatment that was able to achieve functional and histological improvements [48].

This study stands out for showing that the combination did not produce synergistic effects. It is crucial to consider publication bias, which tends to favor the publication of positive results. This bias can create an imbalanced perception of scientific evidence, potentially leading to incorrect conclusions. Publication bias may result in wasted resources and overlook valuable lessons from unsuccessful experiments. Therefore, we emphasize the significance of publishing data, such as the findings presented in this manuscript.

## 4. Materials and Methods

This study has 2 parts, in which Roflumilast was combined with 2 different therapeutic approaches: Levetiracetam and the secretome derived from ASCs. The assessment focused on functional recovery in a rat model of SCI. A total of 72 animals were subjected to a SCI and randomly treated with the therapeutic agents administered alone or given in combination. Functional recovery was assessed during 8 weeks post-injury. All procedures were carried out by EU directive 2010/63/EU and were approved by the ethical committee in life and health sciences (Ref: DGAV 022405, SECVS116/2016, University of Minho, Braga, Portugal). The ARRIVE guidelines for reporting animal research have been followed [49].

### 4.1. Cell culture and secretome collection

Adipose Stem Cells (ASCs), were collected from human donors, according to the procedure described by Dubois [50]. ASCs in passage 6 (P6) were seeded in α-MEM medium (Invitrogen, USA) supplemented with with 5% human platelet lysate (hPL, Stemmatters, Portugal) and 1% Penicillin-Streptomycin (P/S, Invitrogen USA). ASCs were maintained at 37°C and 5% CO2, with medium changes every 2 or 3 days, until reaching confluency. The secretome was collected as described before by our group [51]. Briefly, P6-P8 hASCs were plated at a density of 4 to 12×103 cells/cm2 and left for 72h in a complete α-MEM medium and incubated as previously described. Afterward, the medium was removed, and cells were washed with phosphate-buffered saline (PBS), with no Ca2+ and Mg2+ (Invitrogen, USA). After washing, Neurobasal A medium (NbA, Invitrogen, USA), supplemented with only 1% kanamycin (Invitrogen, USA), was added to the cells. This media was left for 24h at the same incubation parameters. Following this conditioning period, the medium containing the factors/vesicles released by the ASCs – secretome or conditioned media – was collected and centrifuged at 1000g for 10min to pellet cell debris. As a control group we followed the same parameters but without cells in the culture flasks. The secretome or Neurobasal medium only were stored at -80°C, after a snap freeze in liquid nitrogen, until it was required for further experiments.

### 4.2. In vivo study design

The objective of this study was to evaluate the therapeutic efficacy of two distinct treatment combinations, specifically the synergistic effects of Lev and Rof in the first segment, and the complementary interplay between Rof and ASCs secretome (Sec) in the second segment. The assessment focused on the enhancement of functional recovery in a rat model of SCI. In the initial part of the study, a total of 48 animals were allocated into four groups, each consisting of 12 rats. These groups received treatments with Rof alone, Lev alone, a combination of Rof and Lev, or a control group administered with saline. In the latter portion of the study, a total of 24 animals were similarly divided into four groups, with six rats per group. These groups were treated with Rof alone, Sec alone, a combination of Rof and Sec, or Neurobasal alone, serving as the control group. All data collection procedures were conducted in a double-blind manner to minimize any potential bias related to the treatment groups. All procedures were carried out by EU directive 2010/63/EU and were approved by the ethical committee in life and health sciences (Ref: DGAV 022405, SECVS116/2016, University of Minho, Braga, Portugal).

### 4.3. Spinal Cord Injury Model and Treatment

Wistar Han female rats (8 to 10 weeks old, weighing 210–260 g) were used in this study. Animals were kept under standard laboratory conditions (12 hours light: 12 hours dark cycles, 22°C, relative humidity of 55%, ad libitum access to standard food and water), and housed in pairs. Animal handling was carried out 3 days before surgery and then subjected to a severe contusive SCI as previously described [48, 52]. Briefly, general anesthesia was induced by intraperitoneal injection (IP) of ketamine (100 mg/mL, Imalgene/Merial, Duluth, GA, USA) and medetomidine hydrochloride (1 mg/mL, Dormitor/Pfizer, New York, NY, USA) mixture, at a volume ratio of 1.5:1. Once anesthetized, animals received subcutaneous injections of the analgesic butorphanol (10 mg/mL, Butomidor/Richter Pharma AG, Wels, Austria), and the antibiotic enrofloxacin (5 mg/mL, Baytril/Bayer, Leverkusen, Germany). The fur was shaved from the surgical site and the skin was disinfected with 70% ethanol and chlorohexidine. Surgical procedures were performed under sterile conditions. The animals were placed in a prone position and a dorsal midline incision was made at the level of the thoracic spine (T5–T12). The paravertebral muscles were retracted and the spinous processes and laminar arc of T8 were removed, exposing the spinal cord. The dura was left intact. A weight drop trauma model was used, which consisted of dropping a 10 g weight rod from a 10 cm height onto the exposed spinal cord. The rod was guided through a stabilized tube that was positioned perpendicularly to the center of the spinal cord. After the trauma, the muscles were sutured with Vicryl suture (Johnson and Johnson, New Brunswick, NJ, USA), and the incision closed with surgical staples (Fine Science Tools, Heidelberg, Germany). Anesthesia was reverted using atipamezole (5 mg/mL, Antisedan/Pfizer, New York, NY, USA).

Post-operative care for all rats included butorphanol (Richter Pharma AG, Wels, Austria) administration twice a day, for five days as well as vitamins (Duphalyte, Pfizer, New York, NY, USA), saline, and enrofloxacin (Bayer, Leverkusen, Germany), twice a day for 7 days. Manual expression of bladders was performed twice a day until animals recovered spontaneous voiding. Body weight was monitored weekly as a parameter of the general health of the animals. If a weight loss over 10% of body weight was detected, a high-calorie oral supplement (Nutri-Cal®) was administered daily.

### 4.4. Combination of Roflumilast and Levetiracetam

For every group, Rof, Lev, Rof + Lev, or Saline administration, an intrathecal catheter was inserted 3 days before inflicting the contusion. Under general anesthesia, a dorsal midline incision was made. The spinous process and laminar arc of T13 were removed, the dura was gently opened, and the catheter was introduced up to the level of T9. Animals were functionally evaluated [Basso, Beattie, and Bresnahan (BBB) scale], at day 2 post-procedure, and were excluded if any motor disability was detected. Three days after catheter implantation the SCI was inflicted as described above, then Lev (750 mg/kg) was administered by IP4 hours after injury, and treatment with Rof was administered locally through the intrathecal catheter (100 µg/kg) on days 2, 4, and 6 post-injury and then weekly by an IP (1 mg/kg) until the eighth week (Figure 5A). This treatment regimen for Rof, where local and systemic administrations were combined, was chosen due to previous positive results [11]. For the Lev + Rof group, both drugs were administered at the same time points and with the same methods of administration for each drug alone. The control group received a saline solution via the same methods and at the same timings as both drugs, while the Lev group also received a local saline solution at the same time points that the Rof group received its treatment and vice-versa.

**Figure 5.**
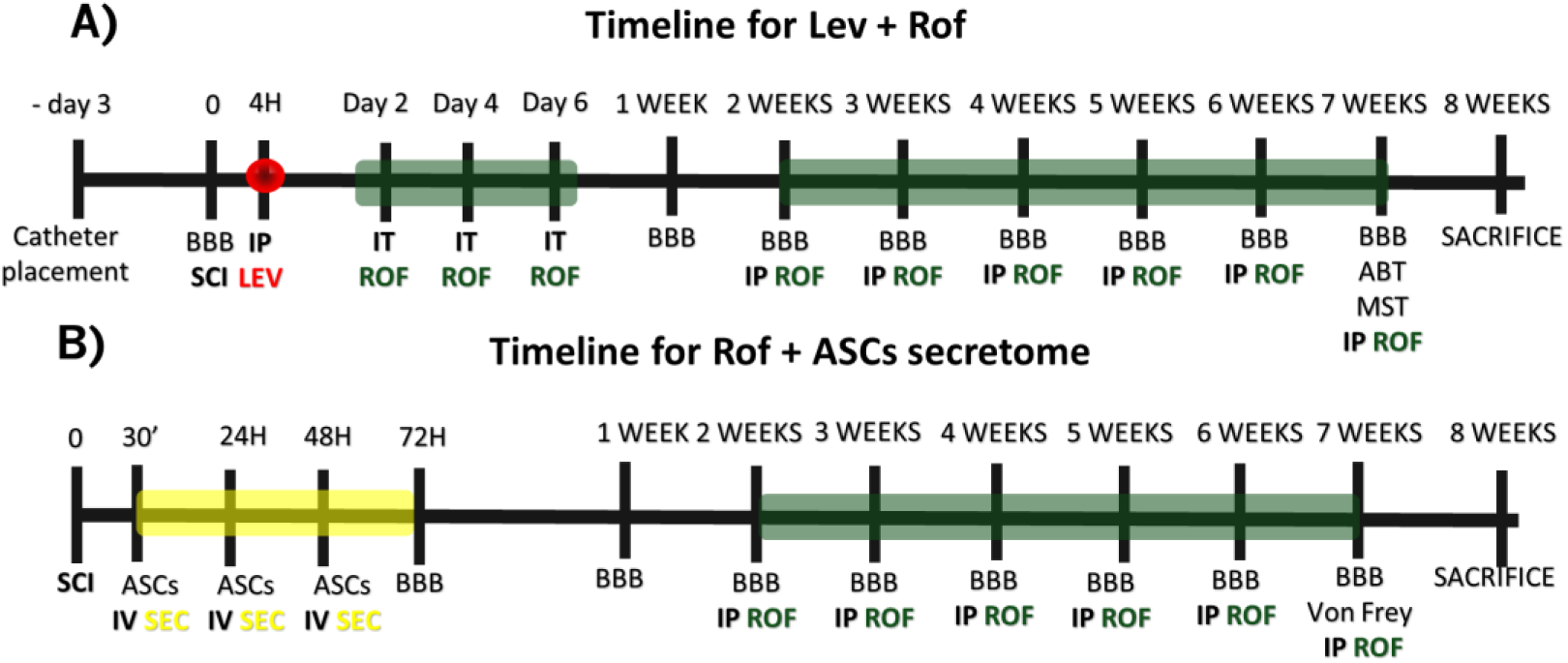
Schematic experimental design. **A**) Therapeutic synergies between Roflumilast (ROF) and Levetiracetam (LEV). Wistar rats underwent intrathecal catheter placement 3 days before spinal cord injury (SCI). LEV was administered by intraperitoneal injection (IP) 4 hours after injury. ROF treatment was administered locally through the intrathecal catheter on days 2, 4, and 6 of pos-injury and then weekly as highlighted in green. Behavioral assessment was performed through Basso, Beattie, and Bresnahan (BBB) scale, Activity Box Test (ABT), Motor Swimming Test (MST) at the timepoints indicated above. **B**) Therapeutic synergies between Roflumilast (ROF) and secretome of ASCs. Wistar rats undergo secretome administration through intravenous injection 30 minutes after SCI, followed by daily administration until 72 hours post-injury, as highlighted in yellow. Treatment with Rof was administered from week 2 and every week until the seventh week by IP as highlighted in green. Behavioral assessment was performed BBB scale and Von Frey test at the timepoints indicated above.

### 4.5. Combination of Roflumilast and ASCs secretome

The secretome of ASCs was administered by intravenous injection (IV) at day 0 (30 min post-SCI), 1, 2, and 3 days post-SCI as previously described 25 and treatment with Rof (1 mg/kg) was administered from week 2 and every week until the seventh week by IP injection (Figure 5B). For the Sec + Rof group, we administered the drugs at the same individual time points and method of administration for each drug alone. The control group received the vehicle solution for each specific treatment, the vehicle for Sec was Neurobasal medium and the vehicle for Rof was saline. Animals treated only with Sec receive saline from week 2 and every week until the seventh week by IP injection. Animals treated only with Rof were administered with Neurobasal medium through (IV) at day 0 (30 min post-SCI), 1, 2, and 3 days post-SCI.

### 4.6. Behavioral Assessment

#### 4.6.1. Basso, Beattie, and Bresnahan (BBB) scale

The BBB locomotor rating scale was used to evaluate functional recovery as previously described 22. The BBB test was executed three days post-injury (dpi) and thereafter weekly for 8 weeks. A BBB score of 0 indicates no hindlimb movement. A BBB score of 1 through 8 indicates joint movement, but no weight support. A BBB score of 9 through 20 indicates an ability to support weight and use the limb for locomotion but with some degree of abnormality. A BBB score of 21 corresponds to the locomotion of a normal rat. All behavioral analyses were executed blindly for the treatment groups.

#### 4.6.2. Activity Box Test (ABT)

This test allows the assessment of exploratory behavior by measuring the total distance traveled, and the velocity of the animals. The test was done in an open arena (43.2 × 43.2 cm) with transparent acrylic walls (Med Associates Inc., Vermont, USA). Animals started the test at the arena’s center and were given 5 minutes to explore it. Data was collected using the activity monitor software. Velocity was used as a measure of locomotor activity [53, 54].

#### 4.6.3. Motor Swimming Test (MST)

As rats swim, they can perform locomotor movements without fully supporting their body weight since water provides buoyancy. When the animals have been trained, they swim without any apparent aversions, and stress symptoms do not significantly affect the test. The motor swimming test was performed on a quadrangular pool (water temperature 25°C), where the rat had to reach the platform to get out of the water. As a training period, the animals performed four trials two days before the test. On the day of behavior analysis, all trials were recorded by a video camera. Velocity was assessed by using three trials of each rat. The software Ethovision XT 12 (Noldus, Wageningen, Netherlands) was used to determine the velocity of swimming.

#### 4.6.4. Von Frey test

The Von Frey test was performed in the seventh week post-injury, as previously described [55]. After 5 minutes of acclimatization to the experimental conditions, animals were placed on an elevated grid. A mechanical allodynia assessment was then conducted using an up-and-down method as described previously [56, 57]. A series of Von Frey calibrated monofilaments: 15.0, 8.0, 6.0, 4.0, 2.0, 1.0, 0.6, and 0.4 g (North Coast Medical Inc., USA) was used to probe the sural dermatome of the lesioned hindlimbs.

Using at first the 2.0 g filament, the test would advance upward if no response was elicited (=0) or downward if a brisk withdrawal of the limb was generated (=X) until 6 measurements were obtained around the threshold point [57]. A response did not include movements of the paws associated with locomotion or weight shifting. The 50% response threshold was then calculated using the following formula: 50%g_threshold=(10Xf+K.δ)/10000.

Where Xf = value (in log units) of the final von Frey filament; k = tabular value corresponding to the pattern of positive and negative responses [X and 0 sequences; consult [56];δ = mean difference (in log units) between stimuli (0.224). If no response was obtained up to maximal force (15.0 g) or conversely, if all filaments elicited a response down to the minimal force (0.4 g), the values 15 and 0.25 were assumed as the 50% withdrawal threshold, respectively.

### 4.7. Statistical Analysis

Statistical analysis was performed using GraphPad Prism 8.00 software. The normality of the data was evaluated by the Kolmogorov-Smirnov normality tests. When the equal variances criterion was not met, Welch correction was applied. Data from the BBB test was assessed by a two-way ANOVA test. Differences between groups were compared with the post hoc Tukey test. Activity box test, motor swimming test, and the Von Frey were analyzed using the non-parametric Kruskal-Wallis followed by the Dunn’s multiple comparisons test. Statistical significance was defined for p < 0.05 (95% confidence level). Data are shown as mean +/− standard error (SEM).

## 5. Conclusions

In conclusion, our study evaluated combinations of Rof with two adjuvant compounds, Lev and the secretome from ASCs, aiming to improve functional recovery after SCI. While the individual treatments with Rof and Lev separately showed efficacy in promoting functional recovery in an SCI rat model, the combined treatment of Rof + Lev did not lead to synergistic effects. However, the combination of Rof + Lev was still significantly better than the vehicle treatment. Contrarily, the combination of Rof with ASCs secretome did not just yield improved functional recovery compared to treatments administered individually but led to detrimental effects. These findings suggest that the combinations tested did not produce superior functional outcomes compared to the standalone treatments. Further investigations are warranted to identify more effective combinatorial treatments with potential clinical applications for SCI.

## Author Contributions

Conceptualization, Nuno Silva; Formal analysis, Carla Sousa, Rui Lima and Eduardo Gomes; Funding acquisition, António Salgado and Nuno Silva; Investigation, Carla Sousa, Rui Lima, Eduardo Gomes, Deolinda Silva, Jorge Cibrão, Tiffany Pinho, Diogo Santos, João Afonso, Joana Martins-Macedo and Andreia Monteiro; Methodology, Rui Lima and Eduardo Gomes; Project administration, Nuno Silva; Supervision, Nuno Silva; Writing – original draft, Carla Sousa; Writing – review & editing, Eduardo Gomes, Andreia Monteiro, António Salgado and Nuno Silva.

## Institutional Review Board Statement

The study was conducted in accordance with the Declaration of Helsinki and approved by the Institutional Ethics Committee of University of Minho (Reference SECVS116/2016) as well as by the national authorities “Direção-Geral da Alimentação e Veterinária” (Reference DGAV022405) for studies involving animals.

## Data Availability Statement

The data that support the findings of this study are available from the corresponding author upon request.

## Acknowledgments

We would like to acknowledge the support from Foundation for Science and Technology (FCT) for grating PhD fellowships to Rui Lima, Deolinda Silva, Jorge Cibrão, Tiffany Pinho, Diogo Santos and João Afonso, as well as the CEEC contract to Nuno Silva.

